# Detailed analysis of paternal knockout *Grb10* mice suggests effects on stability of social behavior, rather than social dominance

**DOI:** 10.1101/493692

**Authors:** Kira D. A. Rienecker, Alexander T. Chavasse, Kim Moorwood, Andrew Ward, Anthony R. Isles

## Abstract

Imprinted genes are highly expressed in monoaminergic regions of the midbrain and their functions in this area are thought to have an impact on mammalian social behaviors. One such imprinted gene is *Grb10*, of which the paternal allele is currently recognized as mediating social dominance behavior. However, there has been no detailed study of social dominance in *Grb10*^*+/p*^ mice. Moreover, the original study examined tube-test behavior in isolated mice 10 months of age. Isolation testing favors more territorial and aggressive behaviors, and does not address social dominance strategies employed in group housing contexts. Furthermore, isolation stress impacts midbrain function and dominance related behavior, often through alterations in monoaminergic signaling. Thus, we undertook a systematic study of *Grb10*^*+/p*^ social rank and dominance behavior within the cage group, using a number of convergent behavioral tests. We examined both male and female mice to account for sex differences, and tested cohorts aged 2, 6, and 10 months to examine any developments related to age. We found group-housed *Grb10*^*+/p*^ mice do not show evidence of enhanced social dominance, but cages containing *Grb10*^*+/p*^ and wildtype mice lacked the normal correlation between three different measures of social rank. Moreover, a separate study indicated isolation stress induced inconsistent changes in tube test behavior. Taken together, these data suggest future research on *Grb10*^*+/p*^ mice should focus on on the stability of social behaviors, rather than dominance *per se*.

## INTRODUCTION

Imprinted genes are defined by their monoallelic, parent-of-origin dependent expression originating from differential epigenetic marks established in the germline (Ferguson-Smith, 2011). This class of genes is highly expressed in the central nervous system and significantly impacts brain development and adult behaviors (Davies, Dent, McNamara, & Isles, 2015). The paternally expressed copy of the imprinted gene *Grb10* (growth factor receptor bound protein 10) is expressed in the developing and adult brain, and we have previously established a potential link to social dominance in mice with disruption of the paternally inherited allele (*Grb10*^+/p^) (Garfield et al., 2011). Murine *Grb10* is located on proximal chromosome 11 and encodes a cellular adapter protein belonging to the small Grb7/Grb10/Grb14 family (Charalambous et al., 2003; Han, Shen, & Guan, 2001). This protein has an inhibitory effect on signaling through receptor tyrosine kinases, including the insulin receptor (IR) and insulin-like growth factor receptor (IGFR) (Desbuquois, Carré, & Burnol, 2013). Paternal *Grb10* is highly expressed in the midbrain and hindbrain, including regions such as the ventral tegmental area, the substantia nigra pars compacta, the dorsal raphe nucleus, thalamus, and hypothalamus, and is neuron-specific (Garfield, 2007; Garfield et al., 2011).

Male *Grb10*^*+/p*^ mice 10 months of age were previously reported to be significantly less likely to back down in the Lindzey tube test. This correlated with an elevated incidence of facial barbering in cages containing *Grb10*^*+/p*^ mutants (Garfield et al., 2011). Both measures are considered indicators of social dominance (Lindzey, Winston, & Manosevitz, 1961; Strozik & Festing, 1981; Wang, Kessels, & Hu, 2014). However, in the original study tube testing was not conducted within an animal’s normal cage group, and also took place after mice were isolated for an extended period to determine whether the barbering was self-inflicted (Garfield et al., 2011). Social isolation impacts midbrain function and dominance-related behaviors, often through alterations in monoaminergic signaling (Angulo, Printz, Ledoux, & McEwen, 1991; Valzelli & Bernasconi, 1979). In periods of isolation between 14 and 28 days, this includes alterations in tyrosine hydroxylase transcription, and over 3 months this includes changes in epigenetic marks and writer/eraser activity in the midbrain (Angulo et al., 1991; Siuda et al., 2014). Even short periods alter signaling and connectivity. Acute social isolation over 24 hours potentiates synapses onto dopamine neurons in the dorsal raphe nucleus (DRN) and alters their AMPA receptor/ NMDA receptor ratio and subunit composition (Matthews et al., 2016). Furthermore, social rank itself impacts the subjective experience of isolation, as dominant mice are more sensitive to the behavioral effects of manipulating DRN dopaminergic activity through optogenetic activation and inhibition (Matthews et al., 2016).

Here we systematically explore social dominance behavior of *Grb10*^*+/p*^ mice. We used convergent measures to assess dominance behavior in socially housed *Grb10*^*+/p*^ mice, including the stranger- and social-encounter Lindzey tube tests, the urine marking test, and characterization of barbering behavior. Both male and female cohorts were used to test for any sex differences. Also, cohorts at 2, 6, and 10 months of age were tested in a cross-sectional study designed to account for any differences that may develop with age. Given the extensive changes to midbrain synaptic function, monoaminergic signaling, and epigenetic regulation induced by social isolation, we saw a need to determine whether the isolation period from the earlier experiment (Garfield et al., 2011) impacted the tube test phenotype observed in *Grb10*^*+/p*^ mice. We therefore replicated the dominance testing of isolated *Grb10*^*+/p*^ mice 10 months of age to determine whether isolation stress was required to precipitate the phenotype. Our results indicate *Grb10*^*+/p*^ mice are not more dominant, but may show a social instability phenotype.

## MATERIALS AND METHODS

### Animals

All procedures were conducted in accordance with the requirements of the UK Animals (Scientific Procedures) Act 1986, under the remit of Home office license number 30/3375 with ethical approval at Cardiff University. *Grb10* heterozygous knockout mice on a B6CBAF1/J background were previously created as described in Garfield et al (2011) using a LacZ:neomycin gene-trap cassette interrupting exon 7 (Garfield, 2007; Garfield et al., 2011). This mouse colony was derived via embryo transfer from a colony in Bath and maintained on exactly the same mixed genetic background. Specifically, breeding stock was maintained with either a B6CBA F1/crl line from Charles River or with an in house mixed B6CBA F1/crl x B6CBA F1/J background. Experimental animals were generated by crossing wildtype (WT) breeding stock with the desired parent of origin heterozygous *Grb10*^+/-^ animal. Dams were placed in individual housing the week prior to full term. This measure was necessary to aid pre-weaning ear clip identification and genotyping of the behavioral cohorts. Mice were weaned between P19 and P28 and sorted into genotype-balanced social cages of 4 mice: 2 wildtypes, 2 *Grb10*^*+/p*^ for behavioral testing. Male mice were genotyped prior to weaning to enable the cage set-up. Females were weaned prior to genotyping and re-sorted into the appropriate set-up as soon as possible. Where possible, animals of the same birth litter were kept together.

All mice were housed in single-sex, environmentally enriched cages (cardboard tubes, shred-mats, chew sticks) of 1-5 adult mice per cage (except for isolation study detailed below). Cages were kept in a temperature and humidity controlled animal holding room (21 ± 2°C and 50 ± 10% respectively) on a 12-hour light-dark cycle (lights on at 7:00 hours, lights off at 19:00 hours). All subjects had *ad libitum* access to standard rodent laboratory chow and water. Cages were cleaned and changed once a week at a regular time and day of the week for minimal disruption. Cages were not cleaned during multiple day testing of the same dominance test, and were half-cleaned between tube testing and urine marking blocks.

### Behavioral testing

The 2, 6, and 10 month cohorts (but not the isolation cohorts) underwent dominance testing, in order, for: stranger tube test, social tube test, and (males only) urine marking (Figure 1). Behavioral testing was limited to a 4-week window to prevent age overlap with the other cohorts. Mice were handled as little as possible up until one week prior to the start of behavioral testing; then they were handled daily for 5 days before beginning testing. Testing was performed in a quiet room lit by a single indirect lamp bulb between 25 and 60 W. Match and cage numbers included in analysis for each behavioral test are reported in Tables 1, 2, and 3 below. A “match” constitutes a *Grb10*^*+/p*^ vs wildtype encounter.

**Table 1.** Male Matches–*Grb10*^+/p^ vs WT. Matches between male *Grb10*^*+/p*^ and WT mice included in analysis of social dominance testing.

| Age | Stranger Tube Matches | Social Tube Matches | Urine Marking Matches | Social Isolation Matches |
| --- | --- | --- | --- | --- |
| **2 months** | 28 | 56 | 44 | — |
| **6 months** | 23 | 51 | 52 | — |
| **10 months** | 23 | 46 | 46 | 10 |

**Table 2.**
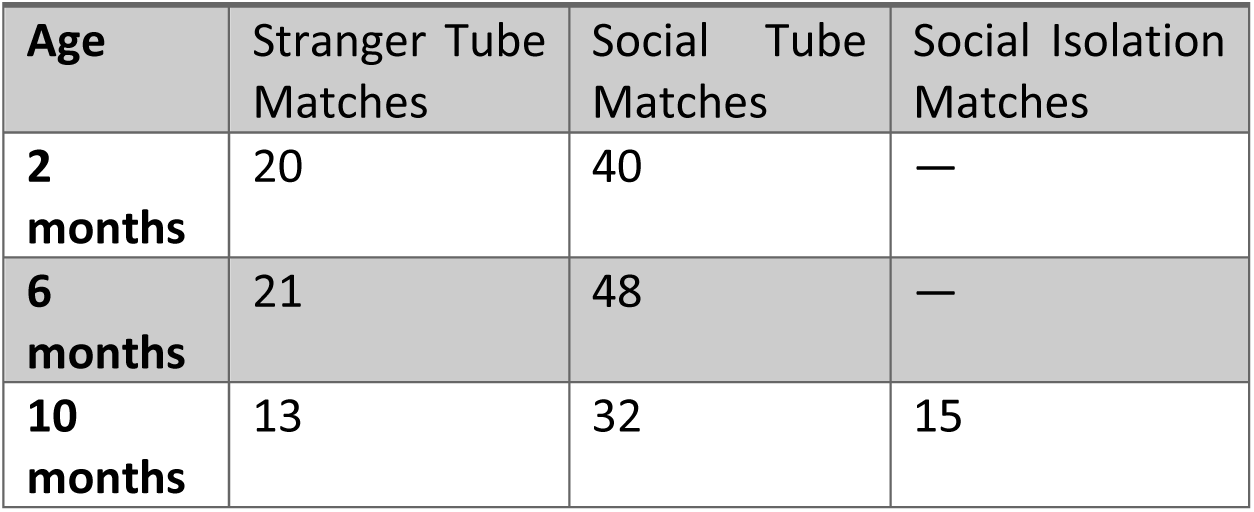
Female Matches– *Grb10*^+/p^ vs WT. Matches between female *Grb10*^*+/p*^ and WT mice included in analysis of social dominance testing.

**Table 3.** Cage Totals in Hierarchy Testing. Cages of mice in each cohort (Male and Female) participating in hierarchy testing.

| Age | Male Cages (Social Tube) | Male Cages (Urine Marking) | Female Cages (Social Tube) |
| --- | --- | --- | --- |
| **2 months** | 15 | 11 | 10 |
| **6 months** | 13 | 13 | 12 |
| **10 months** | 12 | 12 | 8 |

**Figure 1.**
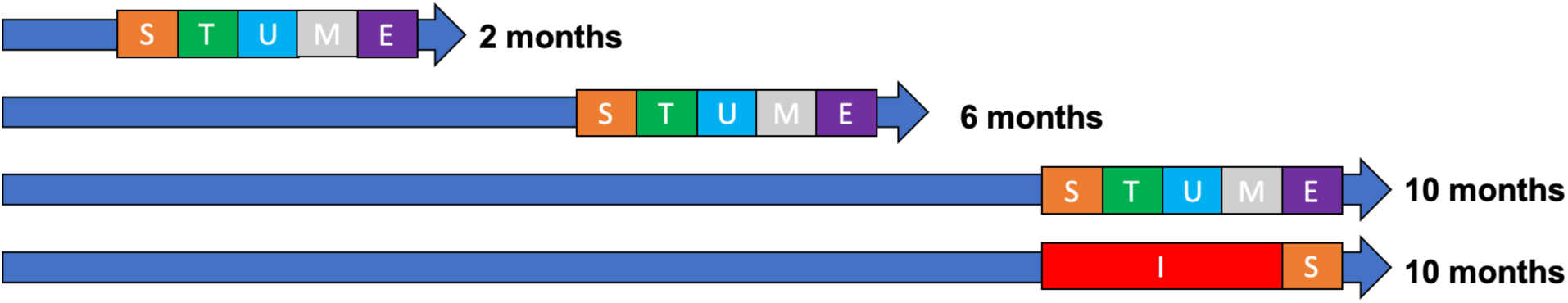
Experimental Design. Four cohorts with both males and females (2 months, 6 months, 10 months, and Isolation cohort at 10 months) underwent behavioral testing. Testing was limited to a 4-week period and ended at the age indicated in the cohort name. The order of experiments was stranger encounter tube test (S), social encounter tube test (T), urine marking test (U; males only), marble burying test (M; not described in this paper), and elevated plus maze (E). The isolation cohort underwent a 30-day isolation protocol (I) prior to the stranger encounter tube test (S).

The Lindzey tube test is an accepted measure of social dominance in mice and can be used to match subjects against strangers or cage-mates (Lindzey et al., 1961). The stranger encounter and social encounter tube tests were conducted under identical conditions. For the stranger test, unfamiliar opponents were chosen from different home cages and different litters. Any socially housed wildtype opponents were housed in genotype-balanced (2 WT, 2 *Grb10*^*+/p*^) cages. Opponent mice were simultaneously presented to either end of a Perspex tube (30.5 m × 3.5cm or 30cm × 2.5cm depending on weight class). Opponents met in the middle of the tube and outcome was scored when one animal was forced to back out. Losers were counted as the first animal with all four feet out of the tube. No time limit was imposed. Trials in which either opponent turned around in the tube, both mice backed out without confrontation, or both mice squeezed past each other were not counted (all instances of trial “failure”). In the stranger encounter tube test, animals were completely naïve to the test and mistrials were not re-run (mistrials are listed in Supplementary Table S1). In the social encounter tube test, mistrials were re-run on a separate day to complete the within-cage hierarchy, but each opponent pair only underwent one successful trial. These paradigms were adopted to avoid any learning effects and to parallel testing procedures in Garfield 2011. Each animal completed only one tube test per day. Testing was arranged to ensure genotype groups and individual mice underwent trials balanced by side of entry. In the stranger encounter tube test, opponents were weight matched to minimize differences across the whole cohort. To maximize trial numbers, no trials were eliminated based on weight. In approximately 77% of encounters, the heavier mouse was less than 15% heavier than the lighter mouse. There were no significant differences in body weight between *Grb10*^*+/p*^ and wildtype mice in our colony across all three ages (2, 6, and 10 months) (See Supplementary Results).

### Urine Marking

Mice were simultaneously placed in one compartment of a 30 × 30 × 30cm box divided by a metal grid. A clear, smooth barrier was placed on top of the grid to prevent escape. Each compartment contained a 14cm by 29.5cm sheet of Whatman chromatography paper (3mm, GE Healthcare UK Limited CAT No 3030-2221). Each trial lasted 1 hour, at the end of which both mice were removed and the cages cleaned with 70% alcohol wipes. Marked paper was stained with Ninhydrin spray reagent (Sigma-Aldrich N1286) and scored using a 1cm^2^ grid overlay. All squares containing a scent mark were counted and used in a ratio against usable grid (total grid squares minus shredded sections and urine marks covering more than 4 consecutive squares). These scent marks/urine drops delineate territorial boundaries and contain chemical cues of social status (Ralls, 1971). The winner of each encounter possessed the higher ratio of squares containing sent marks to usable grid.

### Barbering

The Dhalia Effect, or the whisker barbering effect, describes the tendency for the dominant mouse in the cage to trim the whiskers from subordinates, resulting in cages with just one unbarbered mouse (Strozik & Festing, 1981). Barbering status was recorded at every behavioral testing session. Barbering was identified as the specific removal of whiskers (partial or complete); facial overgrooming could occur independently of barbering, and was thus noted, but not sufficient to confer a ‘barbered’ status.

### Oestrus

Oestrus swabs were taken once per week following behavioral testing. Smears on gelatin-coated slides were stained for 5 minutes using Cresyl fast violet and were identified under the microscope. On other days of testing, a visual assessment of oestrus status was recorded. Statistics pertaining to oestrus use the most closely associated oestrus stage and behavioral testing session.

### Isolation

Socially housed mice 9 months of age were placed in fresh individual housing for 30 days. Immediately following this isolation period, these mice, now 10 months of age, performed the stranger encounter tube test. Mice encountered one unfamiliar mouse of the opposite genotype (*Grb10*^*+/p*^ or wildtype) per day for three days. Cage bedding was not changed during the testing period.

### Statistics

Data analysis was performed using SPSS (versions 23 and 25). Data in diagrams are presented as mean ± standard error of the mean, unless otherwise stated. Statistical significance underwent False Discovery Rate (FDR) corrections using the Benjamini-Liu (BL) method (Y Benjamini & Liu, 1999; Yoav Benjamini, Drai, Elmer, Kafkafi, & Golani, 2001). FDR corrections were performed on all reported measures belonging to one task, and FDR corrections were separate between different tasks. FDR corrections were not carried out for groups of less than 5 statistical tests. The binomial test was conducted to determine if the proportion of *Grb10*^*+/p*^ wins in ‘*Grb10*^*+/p*^*’* versus ‘wildtype’ matches differed significantly from chance (0.5). Most individual mice were involved in two unique matches against cage mates of the opposite genotype. For example, “*Grb10*^*+/p*^ A vs WT B” and “*Grb10*^*+/p*^ A vs WT C” would be included in the analysis as independent matches. The related samples sign test and the Wilcoxon signed-rank test were used to compare the difference in cage rank between the genotype groups. Hierarchies were established in each cage, with rank scored between 0 (least dominant) to 1 (most dominant), based on the number of wins divided by possible matches against cage mates. Data about differences and average genotype rank were presented as medians. The Mantel-Haenszel test of trends was run to determine if there was a linear association between pairs of social tube test rank, urine marking rank, and barbering rank in total male mice at each age cohort. For these statistical analyses, rank was described between 0 (0 wins against cage mates in the dominance tests) and 3 (three wins against cage mates in the dominance tests), or as 0 (barbered subordinate) and 1 (dominant barber).

## RESULTS

### Oestrus and Barbering status did not consistently predict tube test wins

Female mice are commonly excluded from social dominance assessments as they do not share some of the behaviors used to assess male social hierarchies, such as territorial marking and vocalizations to a potential mate. However, female mice can establish stable linear hierarchies in the Lindzey tube test. While test outcomes for male mice are strongly influenced by prior social experience, female mice primarily rely on intrinsic attributes to establish a hierarchy (van den Berg, Lamballais, & Kushner, 2015). Consequently, we tested both male and female mice. Before proceeding with analysis of our *Grb10*^*+/p*^ vs WT matches, we analyzed stranger encounter tube tests in the female cohorts to determine whether we could predict tube test wins using oestrus status. In 16 social tube test matches pooled across the 2, 6, and 10 month cohorts, a wildtype mouse judged to be in oestrus faced a wildtype mouse not in oestrus. A binomial test indicated the proportion of wins for wildtype females in oestrus (0.44) was not significantly different from chance (0.5), p = 0.804 (2-tailed). Further analysis was performed on matches ignoring genotype. In 18 social tube test matches pooled across cohorts, a mouse judged to be in oestrus faced a mouse judged not to be in oestrus. In 9 matches, the mouse in oestrus was *Grb10*^*+/p*^, and in the remaining 9 the mouse in oestrus was wildtype. A binomial test indicated the proportion of wins for mice in oestrus (0.33), regardless of genotype, was not significantly different from chance (0.5), p = 0.238 (2-tailed). Based on these results, we justified ignoring oestrus stage in the statistical analysis of both stranger encounter and social encounter tube tests in the following sections.

We also analyzed the impact of barbering status on the stranger encounter tube test for males and females. In 16 matches in the 6-month cohort, a barbered female mouse faced an unfamiliar, un-barbered female mouse (of a different genotype, as per the design). In 8 matches the barbered mouse was *Grb10*^*+/p*^, and in the remainder, the barbered mouse was wildtype. A binomial test indicated the proportion of wins for barbered female mice 6 months of age (0.88) against unbarbered mice, regardless of genotype, was statistically different from chance (0.5), p = 0.004 (2-tailed). This result survived FDR correction. For males 6 months of age, and males and females 10 months of age, barbering status was unable to predict the outcome of the stranger encounter tube test. No barbering was observed at 2 months. We concluded barbering status did not adequately predict the outcome of a stranger encounter in the Lindzey tube test, and excluded it from our subsequent analyses.

### Grb10^+/p^ barbers were no more common than wildtypes

Garfield 2011 reported an increased incidence of barbering in cages with *Grb10*^*+/p*^ mice. In our study, behavioral cages at 6 months and 10 months with identifiable barbers (1 un-barbered to 3 barbered mice in the cage) were pooled to analyze the proportion of *Grb10*^*+/p*^ vs WT barbers (Table 4). Binomial tests indicated the proportion of barbers who were *Grb10*^*+/p*^ was not statistically different from chance (0.5) in cages of either sex (Figure 1; males p = 0.180, females p = 0.774, two-tailed). After 30 days of isolation, none of the mice showed signs of whisker barbering.

**Table 4.** Barber Genotype in Behavioral Cohorts. Cages with a clear 1:3 dominant barber to subordinate barbered mouse ratio were pooled across age for analysis. There was no statistically significant difference in genotype frequency among barbers.

| Sex | Barbered Cages (pooled) | WT Barbers | *Grb10<sup>+/-</sup>* Barbers | Sig. |
| --- | --- | --- | --- | --- |
| Male | 9 cages | 0.78 | 0.22 | 0.180 |
| Female | 12 cages | 0.58 | 0.42 | 0.774 |

### Socially housed *Grb10*^+/p^ mice do not demonstrate a social dominance phenotype

In the stranger-encounter (Figure 3 A & B; Table 5) and social encounter tube tests (Figure 3 C & E; Table 6 & 7), binomial analysis indicated the proportion of wins for Grb10^+/p^ mice in all three age groups for both sexes were not significantly different to chance (0.5). Likewise, the proportion of *Grb10*^*+/p*^ wins in the urine marking test was not statistically higher than chance in the 6- and 10-month cohorts. In the 2-month cohort, the proportion of *Grb10*^*+/p*^ wins in the urine marking test (0.70) at 2 months of age was statistically higher than chance (0.05), p = 0.01 (2-tailed), but this did not survive FDR corrections (Figure 3D Table 8).

**Table 5.** Stranger Tube Test Grb10^+/p^ Proportions and P-values. Statistical analysis of socially housed *Grb10*^*+/p*^ proportion wins against unfamiliar wildtypes in the stranger encounter tube tests.

| Age | Male matches (N) | Males proportion wins | Males p value | Female matches (N) | Females proportion wins | Females p value |
| --- | --- | --- | --- | --- | --- | --- |
| 2 months | 28 | 0.320 | 0.087 | 20 | 0.350 | 0.263 |
| 6 months | 23 | 0.522 | 1.000 | 21 | 0.619 | 0.383 |
| 10 months | 23 | 0.430 | 0.678 | 13 | 0.462 | 1.000 |

**Table 6.** Males Social Tube Test Grb10^+/p^ vs WT Binomial Analysis. Statistical analysis of socially housed male *Grb10*^*+/p*^ proportion wins in matches against wildtype cage mates in the social tube test and proportion of linear hierarchies in genotype balanced cages completing the social tube test.

| Age | Male matches (N) | Males proportion wins *Grb10 +/p* | Males p value | Linear Hierarchy |
| --- | --- | --- | --- | --- |
| 2 months | 56 | 0.43 | 0.350 | 11/12 |
| 6 months | 51 | 0.51 | 1.000 | 8/12 |
| 10 months | 46 | 0.41 | 0.302 | 9/11 |

**Table 7.**
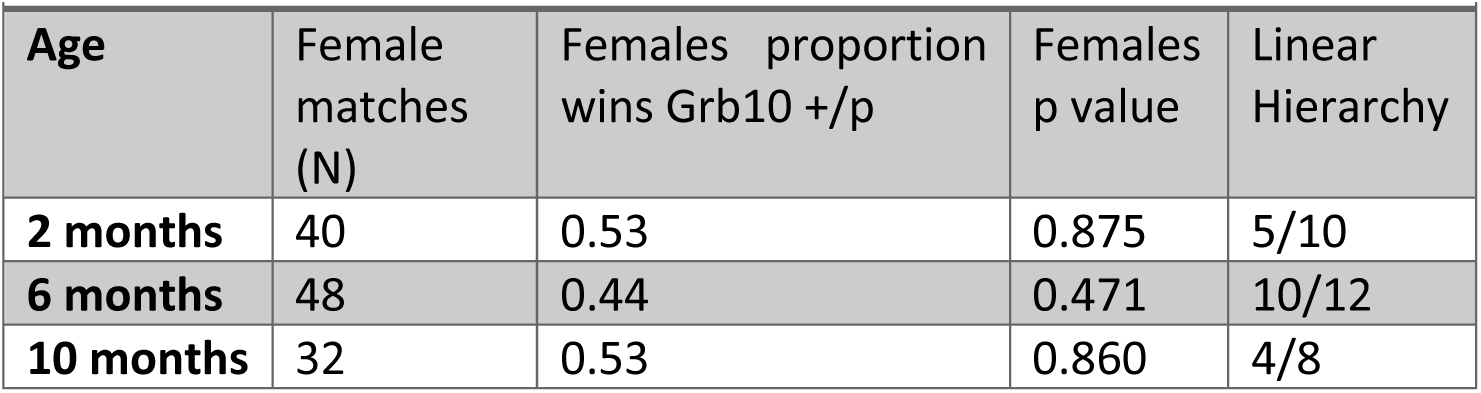
Females Social Tube Test Grb10^+/p^ vs WT Binomial Analysis. Statistical analysis of socially housed female *Grb10*^*+/p*^ proportion wins in matches against wildtype cage mates in the social tube test and proportion of linear hierarchies in genotype balanced cages completing the social tube test.

| Age | Female matches (N) | Females proportion wins $Grb10^{+/p}$ | Females p value | Linear Hierarchy |
| --- | --- | --- | --- | --- |
| 2 months | 40 | 0.53 | 0.875 | 5/10 |
| 6 months | 48 | 0.44 | 0.471 | 10/12 |
| 10 months | 32 | 0.53 | 0.860 | 4/8 |

**Table 8.**
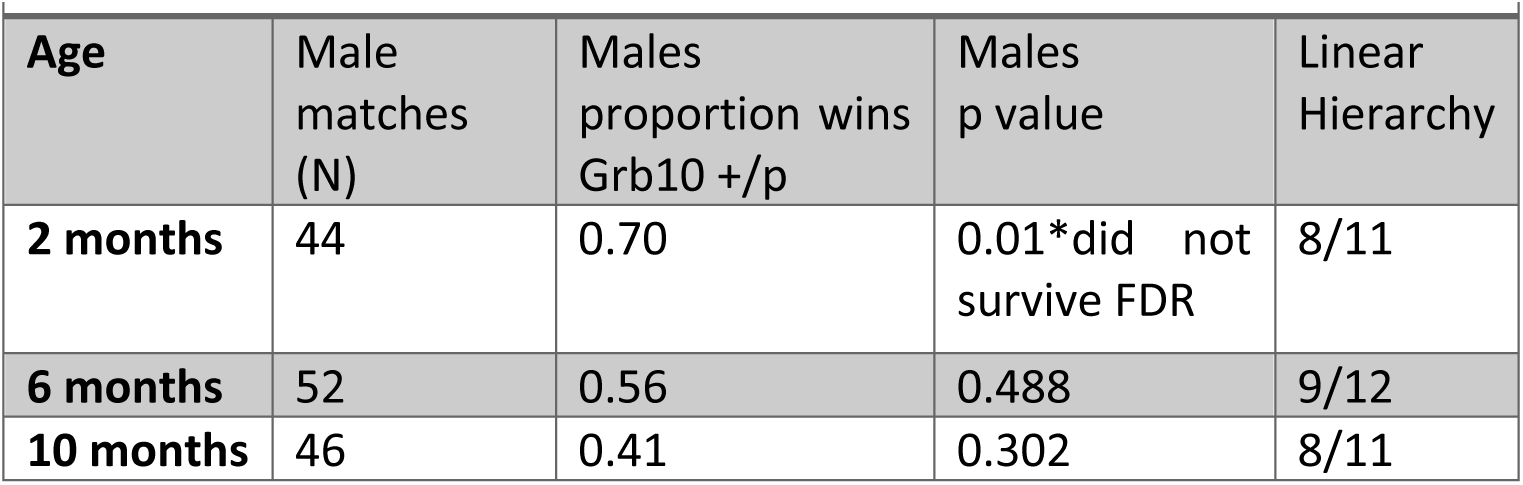
Males Urine Marking Test Grb10^+/p^ vs WT Binomial Analysis. Statistical analysis of socially housed male *Grb10*^*+/p*^ proportion wins in matches against wildtype cage mates in the urine marking test and proportion of linear hierarchies in genotype balanced cages completing the urine marking test. A significant difference at 2 months did not survive FDR correction.

| Age | Male matches (N) | Males proportion wins $Grb10^{+/p}$ | Males p value | Linear Hierarchy |
| --- | --- | --- | --- | --- |
| 2 months | 44 | 0.70 | 0.01*did not survive FDR | 8/11 |
| 6 months | 52 | 0.56 | 0.488 | 9/12 |
| 10 months | 46 | 0.41 | 0.302 | 8/11 |

**Figure 2.**
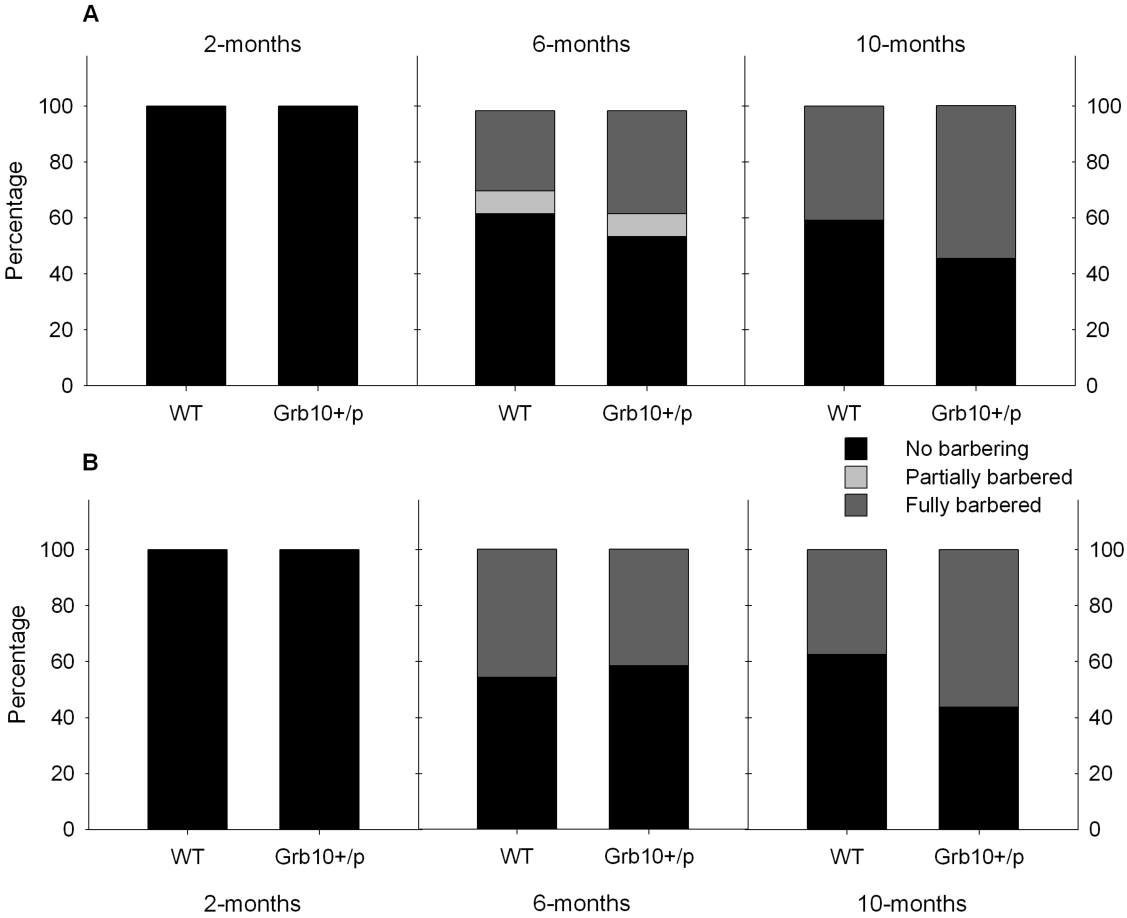
Whisker barbering in *Grb10*^+/p^ socially house mice. Proportions of whisker barbering subdivided by genotype in A) Male and B) Female behavioral cohorts. Barbering was not present at 2 months, but tended to increase with age.

**Figure 3.**
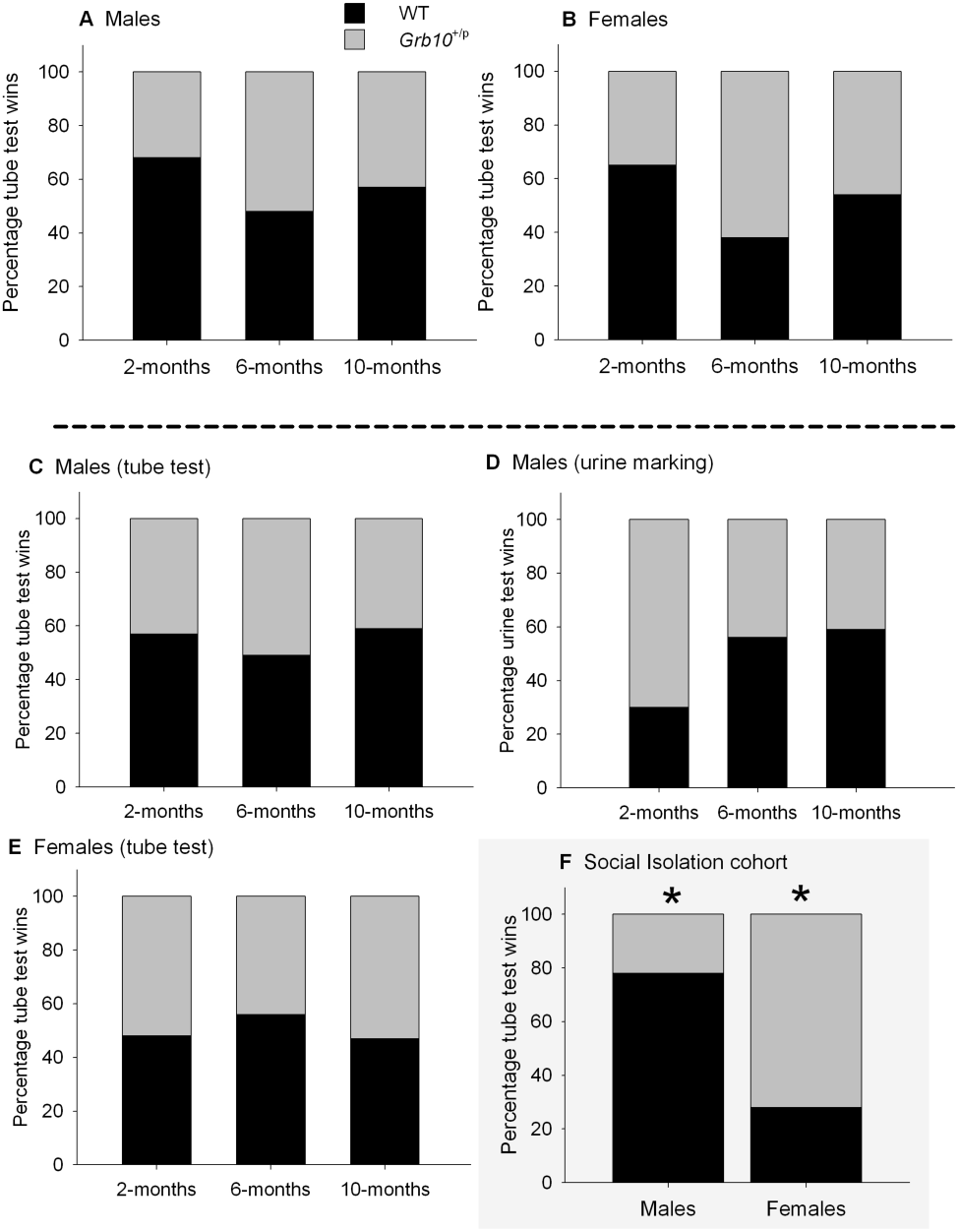
Social dominance tests in *Grb10*^+/p^ mice. A) Male tube test wins vs. strangers. B) Female tube test wins vs. strangers. C) Male tube test wins vs. cage mates. D) Male urine test wins vs. cage mates. E) Female tube test wins vs. cage mates. There were no significant genotype differences for any dominance tests conducted with socially housed mice. F) Male and female tube test wins vs. strangers following 30 days of social isolation.

Rank hierarchies were also established in each cage using the social encounter tube and the urine marking tests (See Supplementary Figure S1). In the social tube test, there was no statistically significant difference between average within-cage rank for *Grb10*^*+/p*^ and wildtypes at 2, 6, or 10 months of age for males or females. In the urine marking test, there was no significant difference in average within-cage rank at 6- and 10 months. At 2 months, the difference in urine marking rank between *Grb10*^*+/p*^ mice (median average cage rank 0.667) and wildtypes (median average cage rank 0.333) was statistically significant, but this did not survive FDR correction. In both Garfield’s testing (light/dark box, open field) and our own (elevated plus maze, see Supplementary Methods, Results and Figure S2), *Grb10*^*+/p*^ mice did not display anxiety phenotypes which might confound social dominance testing (Garfield, 2007; Garfield et al., 2011; Hollis et al., 2015; van der Kooij et al., 2018).

### Social Isolation induces inconsistent effects on *Grb10*^+/p^ dominance behavior

We replicated the social dominance paradigm in Garfield et al (2011) to determine if isolation stress was required to precipitate a social dominance phenotype. Naïve isolated *Grb10*^*+/p*^ mice faced one naïve unfamiliar isolated wildtype per day for three days (Garfield, 2007; Garfield et al., 2011). On Day 1, binomial analysis determined the proportions of male and female *Grb10*^*+/p*^ wins were not statistically significantly different to chance (0.5). Over three days of stranger encounter tube tests, the proportion of male *Grb10*^*+/p*^ wins (0.22) was statistically significantly lower than chance (0.50), p = 0.006 (2-tailed), n = 27 matches. Conversely, the proportion of *Grb10*^*+/p*^ female wins (0.72) over three days of stranger encounter tube tests was significantly higher than chance (0.5), *p* = 0.009 (2-tailed), n = 39 matches (Figure 3F). Both significant results for male and female *Grb10*^*+/p*^ wins over three days survived FDR corrections. Finally, the observed proportion of wins (0.69) for isolated female mice in oestrus, irrespective of genotype, was not statistically different to chance (0.5), *p* = 0.267 (2-tailed), n = 13 matches. Therefore, we also considered isolated males and females together. Over the three-day trial period, the proportion of total *Grb10*^*+/p*^ wins in unique matches (0.52) was not statistically significantly different to chance (0.5), *p* = 0.902 (2-tailed).

### Socially housed mixed genotype cages show signs of social hierarchy instability

While male cohorts had a higher absolute proportion of linear hierarchies in the social dominance tests than females (Supplementary Figure S1), both sexes showed evidence of transitivity within each test. Consequently, cage ranks determined by the social tube test, urine marking test, and barbering status were analyzed for linear correlation. Different tests of social dominance are expected to correlate (Wang et al., 2011), and indeed we have previously seen this in our lab (McNamara, John, & Isles, 2018). However, there was no significant linear association between rank in the social tube test and rank in the urine marking test for the male behavioral cohorts 2 and 6 months of age (Figure 4). At 10 months of age there was a significant linear association between tube test rank and urine marking rank, χ^2^(1) = 7.176, *p* = 0.007, *r* = 0.409, n = 44. When this cohort was broken down by genotype group, a significant linear association was found for male *Grb10*^*+/p*^ (χ^2^(1) = 5.706, *p* = 0.017, *r* = 0.521) mice, but not for wildtypes.

**Figure 4.**
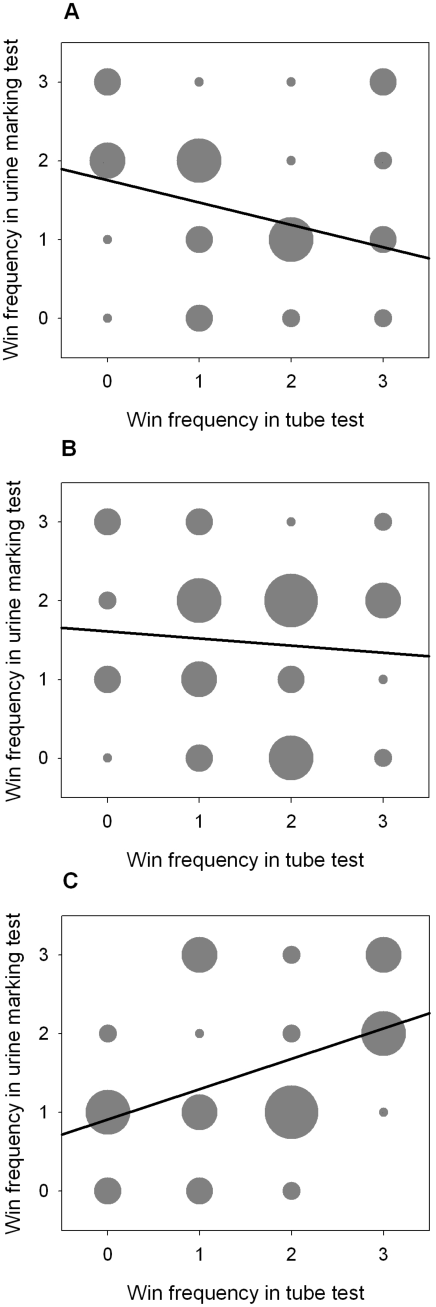
No correlation between social dominance measures in mixed cages of *Grb10*^+/p^ and WT mice. Win frequency (0, 1, 2, or 3 wins) in the urine marking test was plotted against frequency in the social tube test for each male mouse. A) Males 2 months, B) Males 6 months, C) Males 10 months. There was initially a significant linear association at 10 months, but this did not survive FDR correction.

Additionally, there was a significant linear association between tube test and barbering rank for male mice (pooled genotypes) 10 months of age (χ^2^(1) = 3.993, *p* = 0.046, *r* = 0.602, n = 12) (Figure 5). When the cohort was broken down by genotype group, male wildtypes (χ^2^(1) = 4.091, p = 0.043, *r* = 0.905) but not *Grb10*^*+/p*^ mice had a linear association. All other associations between barbering and social tube (male and female) or barbering and urine ranking (male) mice were not significant (Figure 5). Although the four associations of cage rank above were originally found to be significant, none survived FDR correction.

**Figure 5.**
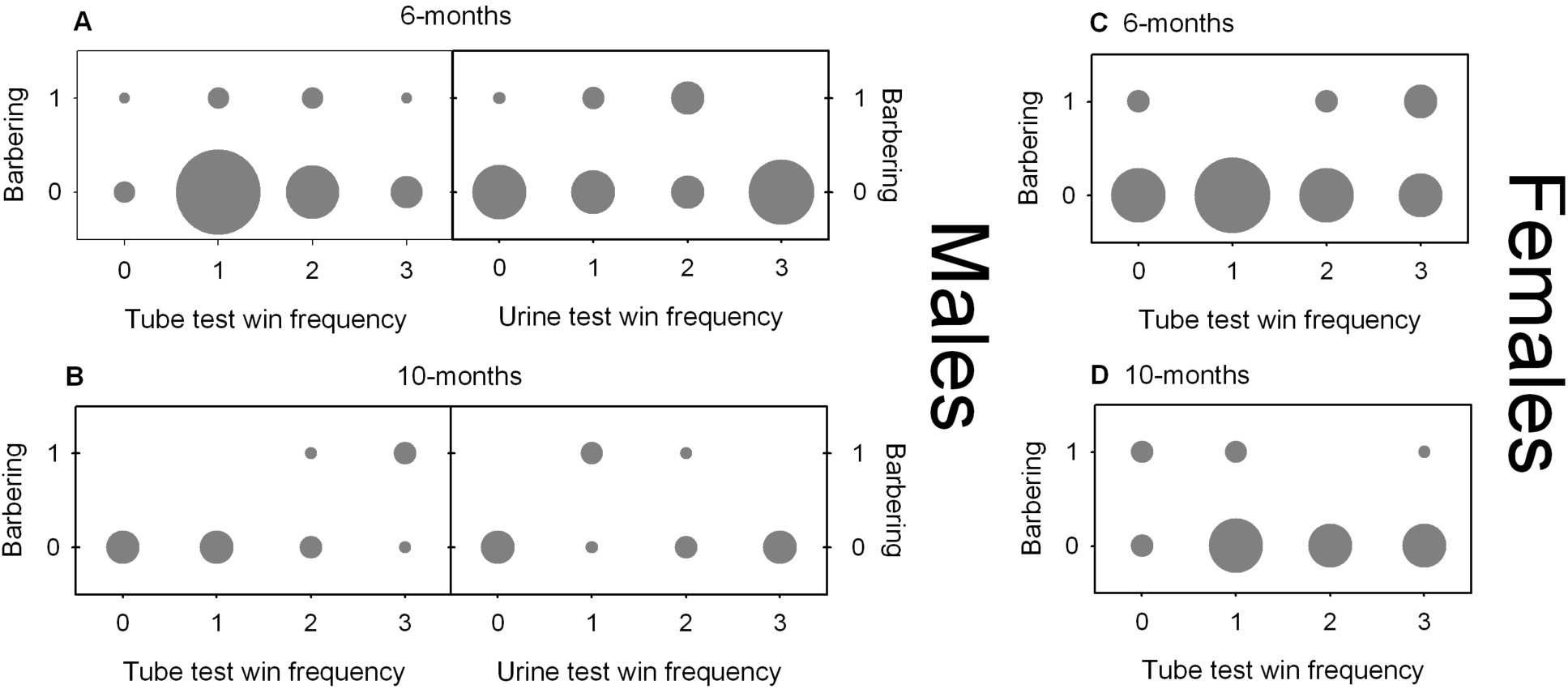
No correlation between social dominance measures and barbering in mixed cages of *Grb10*^+/p^ and WT mice. Barbering status (0–subordinate barbered mouse, 1-dominant barber) was plotted against win frequencies in the social tube and urine tests (0, 1, 2, or 3 wins). Barbering plotted for A) male mice at 6 months against social tube and urine tests, B) male mice at 10 months against social tube and urine tests, C) female mice at 6 months against social tube test, and D) female mice at 10 months against social tube test. There was no barbering at 2 months (See Figure 2).

## DISCUSSION

Our primary goal was to assess social dominance behavior in group-housed *Grb10*^*+/p*^ mice at multiple ages. Social housing provided a more ecologically relevant context for social dominance strategies optimal in close quarters. Group housed animals benefit from social hierarchies reducing costly conflicts, in contrast to isolation housing, where more territorial and aggressive confrontation strategies are more beneficial (Singleton & Krebs, 2007; Wang et al., 2014). We examined three cohorts, at 2, 6, and 10 months of age, to capture any variation in dominance or hierarchical behaviors that might depend on age. Barbering, for instance, was absent in our 2 month cohorts, and appeared in cohorts 6 and 10 months of age. Male and female mice underwent testing to determine whether sex-specific strategies were differentially impacted by paternal *Grb10* deletion (van den Berg et al., 2015).

In both sexes and at all three age groups, we found no difference between *Grb10*^*+/p*^ and wildtype socially housed mice in the likelihood of winning matches in the stranger-encounter Lindzey tube test, familiar-encounter Lindzey tube test, or urine marking test. In the two within-cage dominance tests, we found no significant genotype differences in average cage rank. Additionally, the proportion of *Grb10*^*+/p*^ barbers pooled across all three age groups was not statistically significantly different from chance. From this convergent evidence across large cohorts of both sexes at multiple ages, we concluded socially housed *Grb10*^*+/p*^ mice do not show enhanced social dominance.

These results contrasted the previously reported enhanced dominance phenotype of isolated *Grb10*^*+/p*^ male mice in tube test matches against unfamiliar mice (Garfield et al., 2011). We next replicated the conditions of the Garfield 2011 study to assess whether social isolation stress precipitated the social dominance phenotype in *Grb10*^*+/p*^ mice. In our isolation studies, *Grb10*^*+/p*^ males were statistically significantly *less* likely to win in the stranger-encounter Lindzey tube test against an unfamiliar socially isolated wildtype opponent. This result was opposite to the finding reported in Garfield et al. 2011. Although these experiments were run in different labs (Bath and Cardiff), we replicated the background strain (the mice were derived from the original Bath colony), the conditions of testing, and the power of the experiment (Garfield, 2007; Garfield et al., 2011). Notably, we chose not to use a statistical re-sampling technique such as the Monte Carlo permutation test, due to concerns about amplifying noise (Garfield et al., 2011). In contrast to males, our *Grb10*^*+/p*^ females were statistically significantly *more* likely to win in the stranger-encounter Lindzey tube test.

Our data suggest sex-specific effects of isolation on social dominance behaviors in our *Grb10*^*+/p*^ mice. Sex differences in the expression of (presumably) maternal *Grb10* in muscle have been noted (Welle, Tawil, & Thornton, 2008), but as far as we are aware there are no known sex-differences in terms of paternal *Grb10* expression in the brain (Faisal, Kim, & Kim, 2014), although this has yet to be explored systematically. However, taken together with the Garfield study, the opposing direction of effects in male *Grb10*^*+/p*^ mice following isolation do not suggest enhanced social dominance is necessarily a consistent consequence of social isolation. Rather that there is an interaction between *Grb10* expression and isolation that produces a change in social dominance related behaviors. This may be mediated via altered monoaminergic signaling in the midbrain (Angulo et al., 1991; Valzelli & Bernasconi, 1979). For instance, *Grb10*^*+/p*^ mice lack normal expression in dopamine neurons of the dorsal raphe nucleus (Garfield et al., 2011). This population represents the experience of social isolation, and this experience is modulated by an individual’s prior social rank (Matthews et al., 2016). *Grb10*^*+/p*^ mice possibly experience social isolation stress differently, or employ altered social strategies in hierarchical conflicts following isolation stress (Matthews et al., 2016; Singleton & Krebs, 2007).

Agreement between dominance tests is important in demonstrating a given test measures social dominance as an underlying dependent variable, rather than measuring differences in the sensorimotor skills required to undertake the test. Convergent tests strengthen the description of a robust dominance hierarchy and the characterization of a social dominance phenotype (Wang et al., 2014, 2011). We found both *Grb10*^*+/p*^ male and female cages formed linear hierarchies. We therefore performed tests of rank association between our social tube, urine marking, and barbering data. While four associations were originally significant, none remained so after FDR correction. However, reports of barbering and tube test rank correlations in the literature suggest the use of training prior to the tube test results in correlation between these dominance measures, whereas the absence of training does not result in correlation (Wang et al., 2014). To match the protocols reported in Garfield et al. 2011, and to avoid learning effects, we did not use tube test training, and this may be relevant to interpreting the absence of correlation between barbering and tube test results. Regardless, we note successful correlation between tube test (without training) and urine marking ranks in unrelated control colonies (McNamara et al., 2018).

A comparable phenotype, interpreted as social instability, is present in the *Cdkn1c*^*BACx1*^ mouse model, which overexpresses imprinted cyclin dependent kinase inhibitor 1c (*Cdkn1c*) (McNamara et al., 2018). Social instability has adverse effects on individual fitness including anxiety, stress, and reduced breeding rates (Alexander, 1974; Lardy, Allainé, Bonenfant, & Cohas, 2015; Saavedra-Rodríguez & Feig, 2013). *Cdkn1c*^*BACx1*^ mice do not occupy more dominant ranks than their wildtype cage-mates on any individual measure of within-cage social hierarchy. However, in *Cdkn1c*^*BACx1*^ containing cages, an individual’s rank in one dominance measure did not correlate with its rank in another (McNamara et al., 2018). Clear transitive hierarchies in individual measures of social dominance form in both *Cdkn1c*^*BACx1*^/wildtype and *Grb10*^*+/p*^/wildtype cages, but these are demonstrably unstable in *Cdkn1c*^*BACx1*^ colonies (McNamara et al., 2018). Nevertheless, a different experimental set up is required to determine within-cage rank stability over time for social groups with *Grb10*^*+/p*^ animals. It is also possible *Grb10*^*+/p*^ mice alter the behavior of wildtype littermates, as is the case for *Cdkn1c*^*BACx1*^ and *Nlgn3* (Kalbassi, Bachmann, Cross, Roberton, & Baudouin, 2017; McNamara et al., 2018). Our *Grb10*^*+/p*^ and wildtype balanced cage set up lacks an appropriate independent control group, like cages of *Cdkn1c*^*BAClacZ*^ and wildtype mice (McNamara et al., 2018), to test this.

We have demonstrated through robust and convergent testing at multiple ages, and in both sexes, that socially housed *Grb10*^*+/p*^ mice do not demonstrate a social dominance phenotype. Nevertheless, following social isolation there is an interaction with *Grb10* expression that produces a change in social dominance related behaviors, with a sexually dimorphic direction of effects; critically the direction of effects was contrary to previous findings (Garfield et al. 2011). We also noted an absence of correlation of hierarchical rank between different dominance tests undertaken by *Grb10*^*+/p*^ containing cages, a pattern of behavior previously proposed to indicate instability of social rank (McNamara et al., 2018). Taken together, these findings suggest that paternal *Grb10* may influence stability of social behavior. Nevertheless, although it is clear from the work here and others (Dent et al., 2018) that paternal *Grb10* does impact on brain function generally, further work is required to determine the exact role played in social behavior.

## Supplementary Information

### SUPPLEMENTARY METHODS

**Table S1.** Stranger Encounter Tube Test–Trials Not Counted Due to Failure. Total encounters attempted and failed in the stranger encounter tube test. Only successful trials were included in the analysis.

| Totals | F Attempted | F Failed | M attempted | M Failed |
| --- | --- | --- | --- | --- |
| Cohort D | 24 | 0 | 28 | 0 |
| Cohort C | 24 | 3 | 26 | 3 |
| Cohort A/B | 16 | 3 | 26 | 0 |
| Isolation Day 1 | 15 | 4 | 10 | 2 |
| Isolation Day 2 | 15 | 2 | 10 | 1 |
| Isolation Day 3 | 15 | 0 | 10 | 0 |

#### Elevated Plus Maze (EPM)

The Elevated Plus Maze was carried out in a quiet room with overhead fluorescent lighting, which was necessary for Ethovision detection. The maze consisted of two bisecting white arms 80mm in width by 430mm in length and was elevated 45 cm above the foundation. The opposing pairs of arms were designated “Closed arms” (with 17cm high walls) and “Open arms” (without walls) respectively. The center square of 80mm × 80mm was designated “Middle”. One cage of four mice was carried into the testing room at a time, and remained until all cage mates had individually completed the task. To begin the 5-minute trial, mice were placed in Closed Arm 1. Movement was recorded by the Ethovision detection system, while time for grooming, stretch-attend, and head dips over the edge were scored manually. Between trials, the maze was cleaned with 70% alcohol wipes. Data for Ethovision measures in the EPM task were analyzed using a two-way ANOVA, with AGE and GENOTYPE as between-subjects independent variables, and an Ethovision measure as the dependent variable. Data in main effects analyses are presented as estimated marginal mean ± standard error of the estimated marginal mean, unless otherwise stated. Graphs report descriptive means ± standard error of the descriptive mean, unless otherwise stated. One-way ANOVA was used for each age bin separately when two-way ANOVA was not possible.

### SUPPLEMENTARY RESULTS

**Figure S1.**
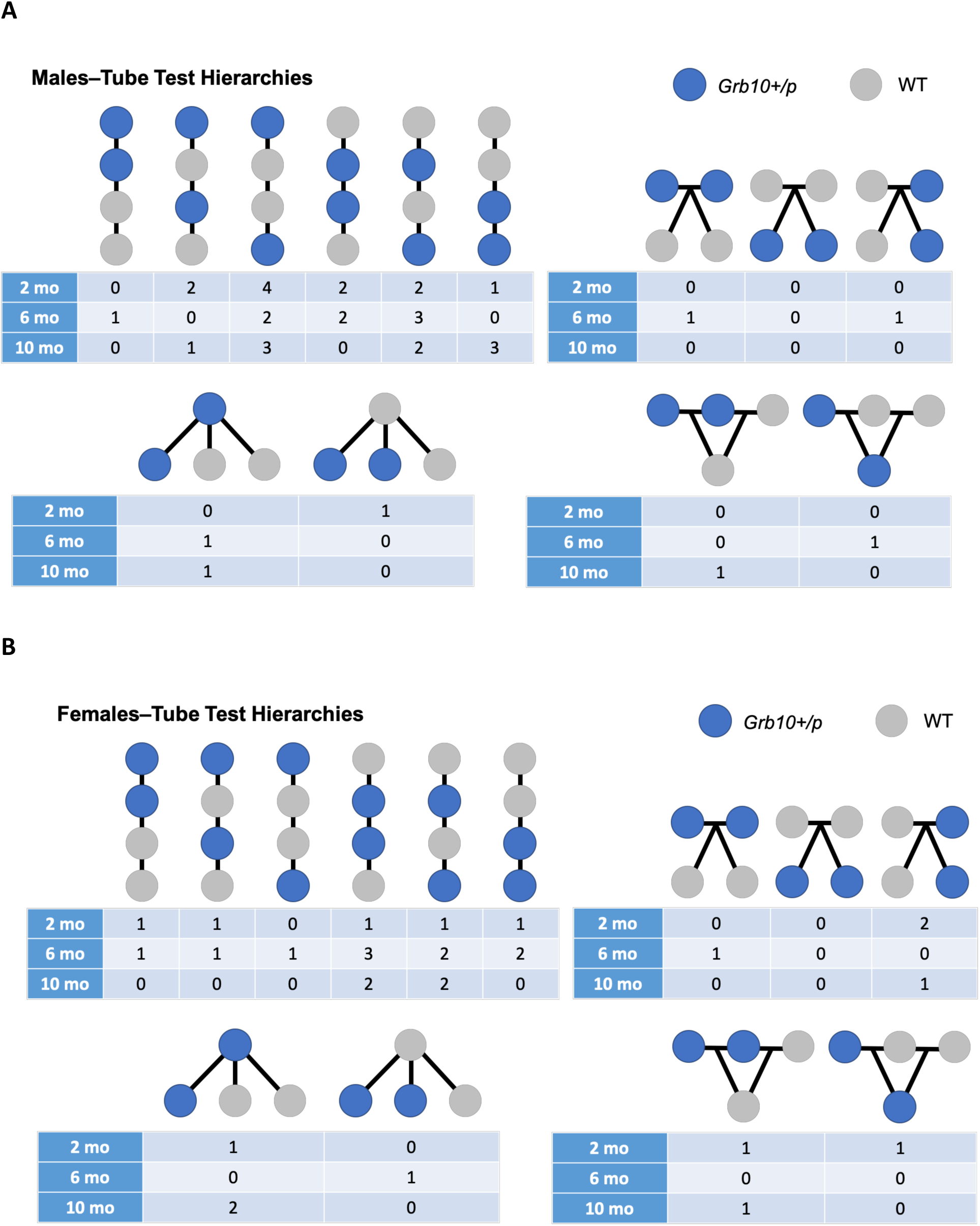
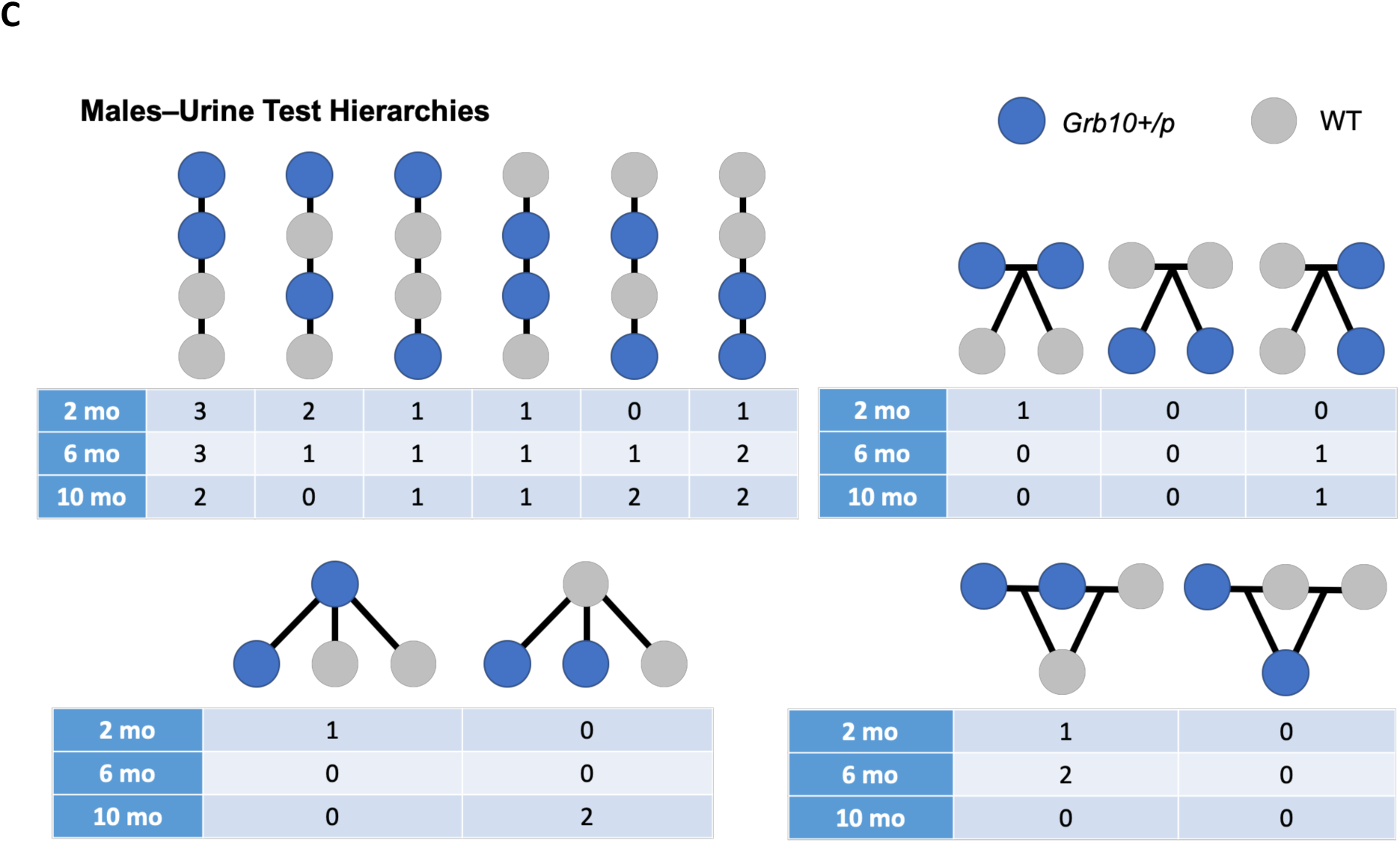
Social dominance hierarchy structures in *Grb10*^+/p^ and WT social groups. All possible hierarchies for 4-mouse cages are depicted with blue (*Grb10*^*+/p*^) and grey (WT) circles connected by lines representing dominance relationships based on number of wins. The most dominant mice (higher number of wins) are towards the top of each diagram. Frequencies of each hierarchy within each cohort are displayed below the diagrams. A) Male social tube test hierarchies, B) Female social tube test hierarchies, C) Male urine marking test hierarchies.

#### Body Weight Analysis

A three-way ANOVA was conducted to examine the effects of between-subjects factors GENOTYPE, AGE, and SEX on body weight in our colony and behavioral groups. GENOTYPE was considered at three levels (wildtype, *Grb10*^*+m*^, and *Grb10*^*+/p*^), AGE was considered at three levels (75-95 days, 185-205 days, and 305-325 days of age), and SEX was considered at two levels (Females and Males). The three-way interaction between GENOTYPE, SEX, and AGE was not statistically significant, nor was the two-way interaction between SEX and AGE. The two-way interactions between GENOTYPE*SEX (F(2,276) = 3.134, p = 0.045, partial η^2^ = 0.022) and AGE*GENOTYPE (F(4,276) = 4.141, p = 0.003, partial η^2^ = 0.057) were statistically significant, and we followed up with simple main effects analysis. For GENOTYPE*SEX, the simple main effect of GENOTYPE was not significant for male or female mice separately, but there was a significant main effect of SEX for wildtype, *Grb10*^*+/m*^, and *Grb10*^*+/p*^ mice individually. Male body weights were consistently heavier than female body weights for all three genotype groups. For AGE*GENOTYPE, the simple main effect of AGE was significant for each genotype group individually. Wildtype and *Grb10*^*+/p*^ mice weighed significantly more at each consecutive age group (75-95 days < 185-205 days < 305-325 days), while *Grb10*^*+/m*^ mice weighed more at 305-325 days than at 185-205 and 75-95 days, but were not significantly different between 75-95 days and 185-205 days. The simple main effect of GENOTYPE for AGE*GENOTYPE was significant for 305-325 days and 75-95 days, but not 185-205 days of age. No pairwise comparisons between the genotype groups at 305-325 days survived Bonferroni correction. At 75-95 days, *Grb10*^*+/m*^ mice were significantly heavier than both wildtype and *Grb10*^*+/p*^ mice, and there was no significant difference between wildtype and *Grb10*^*+/p*^ body weights.

#### Elevated Plus Maze (EPM)

##### Entries to Open Arm

At 10 weeks, total “open arm entries” was not statistically significantly different between *Grb10*^*+/p*^ (19.478 ± 6.626 entries) and wildtype (16.783 ± 7.722 entries) mice, F(1,44) = 1.614, p = 0.211, partial η^2^ = 0.035. At 6 months, “open arm entries” were statistically different between *Grb10*^*+/p*^ (15.700 ± 6.182 entries) and wildtype (9.955 ± 6.484) trials F(1,40) = 8.596, p = 0.006, partial η^2^ = 0.177. This did not survive FDR correction. At 10 months, the assumption of homogeneity of variance was violated (Levene’s test p = 0.019). Therefore, we interpreted Welch’s ANOVA. There was no statistically significant difference in “open arm entries” between *Grb10*^*+/p*^ (16.286 ± 12.546 entries) and wildtype (11.000 ± 6.347 entries) trials, Welch’s F(1,29.300) = 2.995, p = 0.094.

##### Total Entries

As there was a significant genotype difference in total open arm entries at 6 months of age (pre-FDR correction), we also examined total entries to all zones of the EPM to determine if this effect was specific to the open arm. The interaction between GENOTYPE and AGE was not statistically significant for “all entries”, F(2,125) = 0.631, p = 0.534, partial η^2^ = 0.010. Therefore, analyses for main effects were performed. There was a statistically significant main effect of GENOTYPE for “all entries”, F(1,125) = 17.909, p < 0.001, partial η^2^ = 0.125. This survived FDR correction. *Grb10*^*+/p*^ mice made more entries to EPM zones (82.834 ± 2.898 entries) than wildtype mice (65.698 ± 2.828 entries), mean difference 17.137 (95%CI 9.122 to 25.151) entries, p < 0.001.

There was a statistically significant main effect of AGE on “all entries”, F(2,125) = 6.709, p = 0.002, partial η^2^ = 0.097. Mice at 10 weeks made the most entries (84.565 ± 3.413 entries), while mice at 6 months (68.155 ± 3.575 entries) and 10 months (70.078 ± 3.531 entries) made fewer. Mice 10 weeks of age made significantly more entries than mice at 6 months, mean difference 16.411 (95% CI 4.417 to 28.404) entries, p = 0.004. Mice 10 weeks of age also made 14.487 (95%CI 2.572 to 26.402) entries than mice at 10 months, p = 0.011. There was no statistically significant difference between “all entries” made by mice at 6 months and 10 months, mean difference −1.923 (95%CI −14.117 to 10.270) entries, p = 1.000. The main effect of AGE and the pairwise comparisons did not survive FDR correction. Overall, there was a significant genotype difference in “all entries” made to zones of the EPM, indicating increased entries by *Grb10*^*+/p*^ mice at 6 months was not specific to the open arm.

##### Percent Time in Open Arms

As the increase in entries made by *Grb10*^*+/p*^ mice was not specific to the open arm, we examined the division of time to determine whether *Grb10*^*+/p*^ mice differed in the amount of time spent on the open arm. There was no statistically significant interaction between GENOTYPE and AGE for “percent time in open arms”, F(2,125) = 1.226, p = 0.297, partial η^2^ = 0.019. Therefore, analyses for main effects were performed. There was a statistically significant main effect of GENOTYPE on “percent time in open arms”, F(1,125) = 7.727, p = 0.006, partial η^2^ = 0.058. *Grb10*^*+/p*^ mice spent significantly more time on the open arm (19.094 ± 1.390%) than wildtypes (13.697 ± 1.356%), mean difference 5.398 (95%CI 1.555 to 9.241) %, p = 0.006. This effect of GENOTYPE did not survive FDR correction.

There was a statistically significant main effect of AGE on “percent time in open arms”, F(2,125) = 5.786, p = 0.004, partial η^2^ = 0.085. Mice 10 weeks of age spent 20.823 ± 1.636%, 6 months of age spent 15.289 ± 1.715%, and 10 months of age spent 13.074 ± 1.693% of the total time on open arms. Time at 10 weeks was statistically higher than at 10 months (7.749 (95%CI 2.035 to 13.462) %, p = 0.004, but not than at 6 months (5.534 (95%CI −0.217 to 11.285) %, p = 0.063. There was no significant difference between percent time spent on open arms at 6 months and 10 months (2.214 (95%CI −3.633 to 8.062) %, p = 1.000. Neither the main effect of AGE, nor the pairwise comparisons survived FDR correction.

*Grb10*^*+/p*^ mice make more total entries to EPM zones than wildtypes, but do not make more entries to the open arm, nor spend more time on the open arm, when analyses are adjusted for FDR.

**Figure S2.**
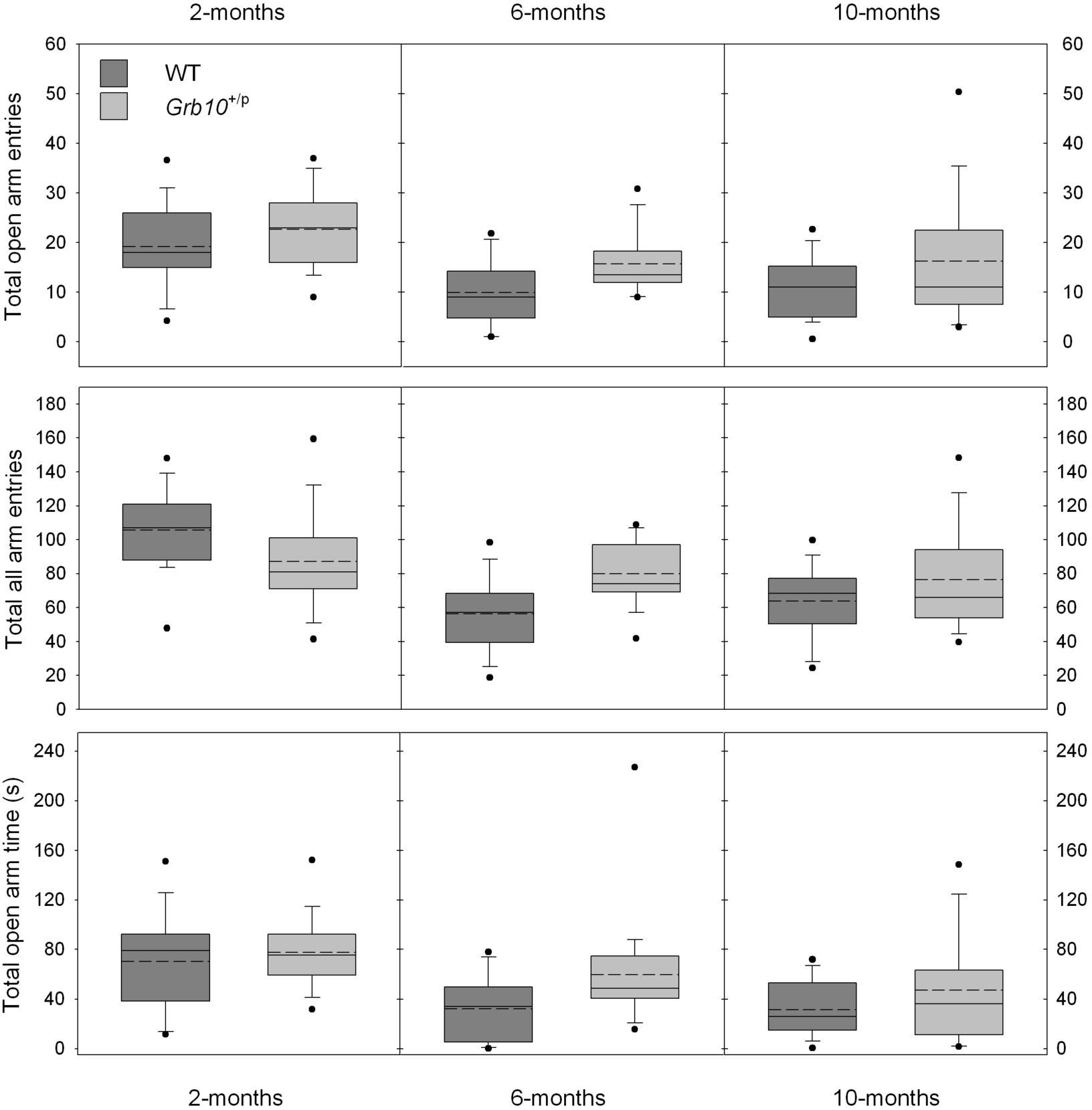
Behavior of *Grb10*^+/p^ mice in the elevated plus maze. Performance in the EPM for 2 month, 6 month, and 10 month male behavioral cohorts. Male *Grb10*^*+/p*^ mice made significantly more total all arm entries, but did not make more total open arm entries (after FDR correction) or spend more total time on the open arm. Data are box-plots showing median (solid line), mean (dashed line) and 5^th^, 25^th^ and 75^th^ and 95^th^ percentiles

